# Opposing action of photosystem II assembly factors RBD1 and HCF136 underlies light-regulated *psbA* translation in plant chloroplasts

**DOI:** 10.1101/2024.11.14.623664

**Authors:** Margarita Rojas, Prakitchai Chotewutmontri, Rosalind Williams-Carrier, Susan Belcher, Emily Boyce, Alice Barkan

**Author notes:** Equal contributions (Margarita Rojas, Prakitchai Chotewutmontri). Crop Improvement and Genetics Research, Western Regional Research Center, United States Department of Agriculture—Agricultural Research Service, Albany, CA 94710.

## Abstract

The D1 reaction center protein of photosystem II (PSII) is subject to light-induced damage. In plants, D1 photodamage activates translation of the chloroplast *psbA* mRNA encoding D1, providing D1 for PSII repair. Genetic data have implicated three D1 assembly factors in the regulatory mechanism: HCF244 and RBD1 activate *psbA* translation whereas HCF136 represses *psbA* translation in the dark. To clarify the regulatory circuit, we analyzed *psbA* ribosome occupancy in dark-adapted and illuminated *rbd1* and *hcf136;rbd1* double mutants in Arabidopsis, and in Zm-*hcf244* and Zm-*hcf136;*Zm-*hcf244* double mutants in maize. The results show that RBD1 is required for the light-induced recruitment of ribosomes to *psbA* mRNA but has little effect on *psbA* ribosome occupancy in the dark. Furthermore, RBD1 is dispensable for *psbA* translation when HCF136 is absent, indicating that RBD1 activates *psbA* translation indirectly, by inhibiting HCF136’s repressive effect. By contrast, HCF244 is required to recruit ribosomes to *psbA* mRNA in light, dark, and in the absence of HCF136. These results demonstrate that the step in D1 assembly that is mediated by RBD1 is central to the perception of the D1 photodamage that triggers D1 synthesis, that RBD1 activates *psbA* translation in the light by relieving the repressive effect of an HCF136-dependent assembly intermediate, and that HCF244 acts independently of these events to activate *psbA* translation. The results implicate a feature of nascent D1 that is affected by both HCF136 and RBD1 as the signal that reports D1 photodamage to regulate *psbA* translation rate as needed for PSII repair.

**Significance Statement:** The D1 subunit of photosystem II (PSII) is damaged by light and must be replaced with nascent D1 to maintain photosynthesis. In plants, D1 photodamage triggers D1 synthesis by activating translation of chloroplast *psbA* mRNA encoding D1. Our results demonstrate that an antagonistic interplay between two photosystem II assembly factors, HCF136 and RBD1, is at the core of the signal transduction process that senses light-induced D1 damage to regulate *psbA* translation. These findings implicate a feature of nascent D1 that is affected by both HCF136 and RBD1 as the signal that reports D1 photodamage to tune *psbA* translation as needed for PSII repair. The results elucidate an assembly-coupled translational rheostat that maintains photosynthesis in the face of PSII photodamage.

## INTRODUCTION

Photosystem II (PSII), a large protein-pigment complex found in thylakoid membranes of chloroplasts and cyanobacteria, harnesses the energy of sunlight to extract electrons from water, producing molecular oxygen and fueling downstream steps in photosynthesis (1). PSII assembly is a complex process that is orchestrated by a suite of assembly factors, and involves the step-wise association of several subassemblies (2). The structure of mature PSII is dynamic due to the sensitivity of its reaction center protein D1 to photooxidative damage. D1 photodamage triggers an elaborate repair mechanism involving partial disassembly of PSII, degradation of damaged D1, and its replacement with newly synthesized D1 (3, 4). Reassembly of the reaction center during PSII repair likely employs many of the same proteins that promote the *de novo* assembly of the PSII core (2).

In photosynthetic eucaryotes, the chloroplast genome encodes D1 and other core PSII subunits whereas nuclear genes encode subunits of the peripheral light-harvesting and oxygen-evolving complexes. Under moderate light intensities, D1 is the most rapidly synthesized protein in the leaves of plants. D1 synthesis drops precipitously shortly after shifting plants to the dark, is restored within minutes of reillumination, and increases when plants are shifted from low intensity to high intensity light (5, 6). These effects result from translational control of chloroplast *psbA* mRNA, which encodes D1. Light regulates *psbA* translation via two superimposed mechanisms in plants (6). A plastome-wide change in the rate of translation elongation modulates the synthesis of all plastid-encoded proteins while maintaining ribosome occupancy on all chloroplast mRNAs other than *psbA*. A *psbA*-specific response involves rapid light-induced changes in translation initiation: *psbA* ribosome occupancy increases dramatically soon after a shift to the light and drops quickly after a shift to the dark (6). These two responses are triggered by different signals: the plastome-wide change in translation elongation rate is regulated by a product of photosynthetic electron transport whereas the light-induced recruitment of ribosomes to *psbA* mRNA is induced by D1 photodamage (7). Thus, light-regulated *psbA* translation initiation in mature chloroplasts is an integral component of the PSII repair cycle, coupling D1 synthesis to D1 damage even under low and moderate light intensities.

Genetic data have demonstrated that early steps in the assembly of the PSII reaction center are central to the regulatory mechanism controlling *psbA* translation in response to light and D1 photodamage. HCF136, which is found in the thylakoid lumen and promotes a very early step in D1 assembly (8-10), represses the recruitment of ribosomes to *psbA* mRNA in the dark (7). HCF244, which is bound to the stromal face of the thylakoid membrane and acts after HCF136 to promote the incorporation of D1 into an early PSII assembly intermediate (2), is required for *psbA* translation initiation (11, 12). These and other data led to a model involving an assembly coupled translational autoregulatory circuit that links D1 synthesis to D1 photodamage (7). The likely endpoint of this signal transduction chain is the translational activator HCF173, which binds the *psbA* 5’ UTR and stimulates ribosome recruitment, at least in part, by maintaining a structure-free translation initiation region on the mRNA (13-16).

New aspects of this phenomenon have come into focus since the time the prior assembly-coupled regulatory mechanism was proposed. First, another PSII assembly factor, RBD1, has been shown to activate *psbA* translation (17). RBD1 is an integral thylakoid membrane protein with a stroma-exposed rubredoxin domain, and affects D1 conformation possibly by facilitating the binding of a non-heme iron to the “DE” loop connecting D1’s fourth and fifth transmembrane segments (18, 19). Ribosome profiling and polysome data showed that *psbA* ribosome occupancy is strongly reduced in Arabidopsis *rbd1* mutants harvested in the light (17). Thus, two D1 assembly factors, HCF244 and RBD1, activate *psbA* translation, whereas another, HCF136, represses *psbA* translation specifically in the dark. Very recently, we found that the *psbA* open reading frame (ORF) acts in *cis* to repress translation initiation in the dark, and that HCF173 itself mediates this repression (20). We proposed an updated model that accounts for most of the observations to date by positing that HCF136 promotes co-translational repressive interactions between HCF173 and nascent D1, and that D1 photodamage relieves this repression via effects on RBD1 and/or HCF244 (20).

The goal of this study was to clarify the roles of the PSII assembly factors RBD1 and HCF244 in light-regulated *psbA* translation by addressing their effects on *psbA* ribosome occupancy in dark-adapted plants, and by epistasis analysis using double mutants with *hcf136*. The results showed that RBD1 enhances *psbA* translation in the light but not in the dark, demonstrating RBD1 to be an essential component of the mechanism that senses D1 photodamage to trigger *psbA* translation. Furthermore, analysis of *rbd1;hcf136* double mutants showed that RBD1 activates translation indirectly by opposing HCF136’s repressive effect. By contrast, we found that HCF244 stimulates *psbA* translation in both the light and the dark, and that it is required for *psbA* translation even in the absence of HCF136. These findings clarify the order of action of HCF136, HCF244, and RBD1 in the signal transduction chain underlying light-activated *psbA* translation and elucidate the step in D1 assembly that influences *psbA* translation in response to D1 photodamage.

## RESULTS

### RBD1 is required for light-activated *psbA* translation and has little effect on *psbA* ribosome occupancy in dark-adapted seedlings

A previous ribosome profiling (ribo-seq) analysis demonstrated a large reduction in the average number of ribosomes on *psbA* mRNA in illuminated Arabidopsis *rbd1* mutants (17). To explore the role of RBD1 in the regulation of *psbA* translation in response to light, we used ribo-seq to analyze *rbd1* mutants harvested after one hour of dark adaptation and after 15 minutes of reillumination (Fig. 1). The published data are displayed adjacent to ours to enable a direct comparison (Fig. 1 left). Despite differences in plant age and growth conditions, the normalized abundance of *psbA* ribosome footprints in illuminated *rbd1* mutants and Col-0 controls was similar in the two experiments. Col-0 showed the expected light-induced increase in *psbA* ribosome occupancy (Fig. 1 right). By contrast, *psbA* ribosome occupancy in illuminated *rbd1* mutants was similar to that in dark-adapted Col-0 and was only modestly decreased in the dark. Therefore, RBD1 is required to stimulate the recruitment of ribosomes to *psbA* mRNA in the light but has little effect on *psbA* translation in the dark. We refer to this dysfunction in *psbA* translational regulation in *rbd1* as an “uninducible” phenotype below.

**Figure 1.**
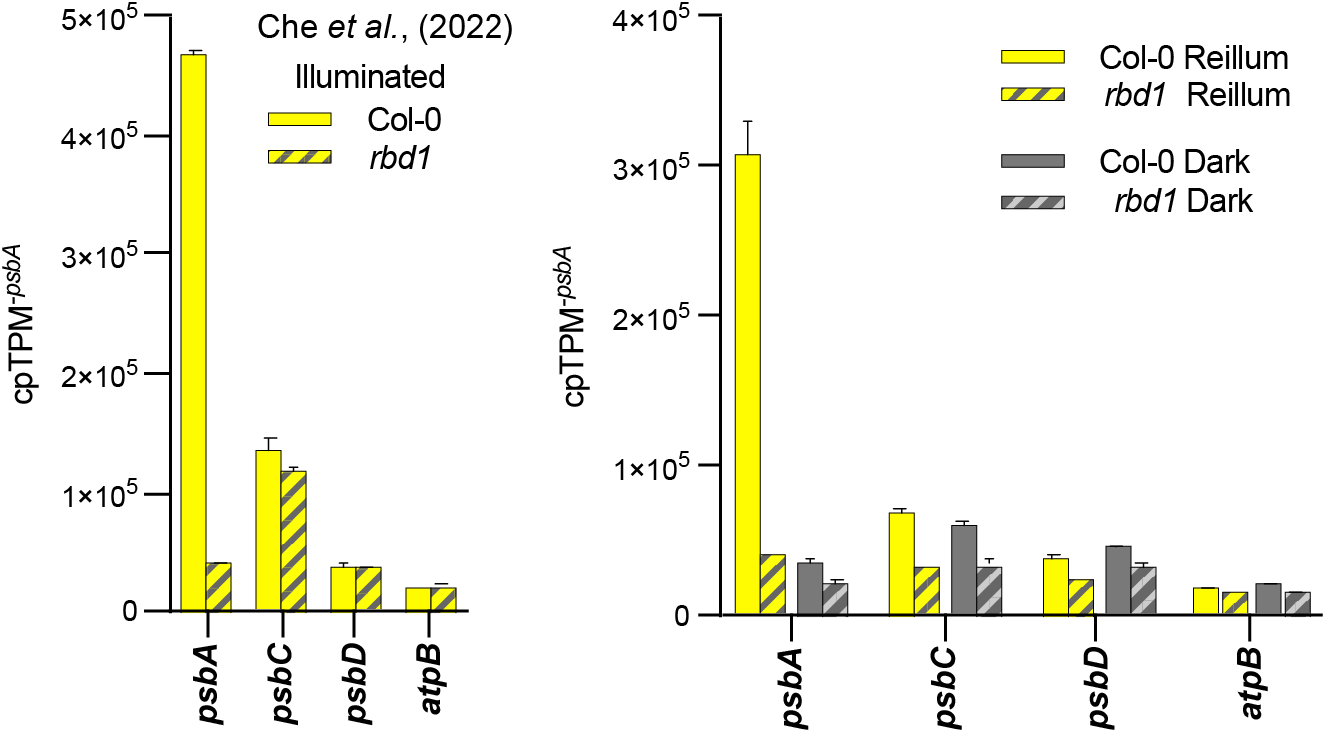
Ribo-seq analysis addressing the effects of RBD1 in dark-adapted and reilluminated Arabidopsis seedlings. The normalized abundance of ribosome footprints mapping to the indicated chloroplast genes in plants harvested after 1-h of dark adaptation or 15-min or reillumination is shown to the right. These data comprise a subset of the data shown in Figure 2, where the abundance of *psbA* mRNA is shown to be similar in the lysates used for ribo-seq. Data were normalized with the cpTPM^-*psbA*^ method as described in Materials and Methods. Published data for illuminated *rbd1* (Che et al., 2022) were normalized using the same method, and are shown to the left for comparison. Values represent the mean +/-SEM of two replicates with the exception of the reilluminated *rbd1* sample, for which the result of a single replicate is corroborated by the results for illuminated *rbd1* reported by Che et al. (left panel). Data for all chloroplast genes are provided in Dataset S1.

### RBD1 has little effect on *psbA* translation in the absence of HCF136

RBD1 and HCF136 both act early in the assembly of D1 into the PSII reaction center (reviewed in 2) and both are essential to regulate *psbA* translation in response to light. However, they affect *psbA* translation in opposite ways. Whereas *psbA* translation in *rbd1* mutants is uninducible (Fig. 1), *psbA* translation in Arabidopsis and maize *hcf136* mutants is “constitutive”: that is, *psbA* ribosome occupancy in the light is normal but it does not drop after shifting plants to the dark (7) (Fig. S1). HCF136 resides in the thylakoid lumen so its effect on translation must be indirect; presumably, HCF136 influences *psbA* translation via the specific way in which it affects D1 assembly. By contrast, RBD1 has a stromal rubredoxin domain (19), so RBD1 could potentially influence translation directly, via interactions with translational regulators, ribosomes, or even *psbA* mRNA. To clarify the roles of RBD1 and HCF136 in the regulatory circuit, we determined whether the constitutively high *psbA* ribosome occupancy observed in *hcf136* mutants requires RBD1. To do so, we analyzed Arabidopsis *rbd1;hcf136* double mutants, harvesting leaf tissue either after 1 hour in the dark or 15 minutes of reillumination.

Progeny of self-pollinated *rbd1/+, hcf136/+* plants were sown on sucrose-containing agar medium, and single and double mutants were identified by PCR analysis of small tissue samples. Homozygous *rbd1* mutants, *hcf136* mutants, and *rbd1;hcf136* double mutants exhibited similar slow growing, pale green phenotypes (Fig. 2A). The abundance of HCF173 and HCF244 was unaffected in *rbd1* and *hcf136* mutants (Fig. 2B), indicating that the effects of RBD1 and HCF136 on *psbA* translation do not result from retrograde effects on the expression of *HCF173* or *HCF244*. The abundance of *psbA* mRNA was similar in all of the genotypes in both the dark-adapted and reilluminated conditions (Fig. 2C).

**Figure 2.**
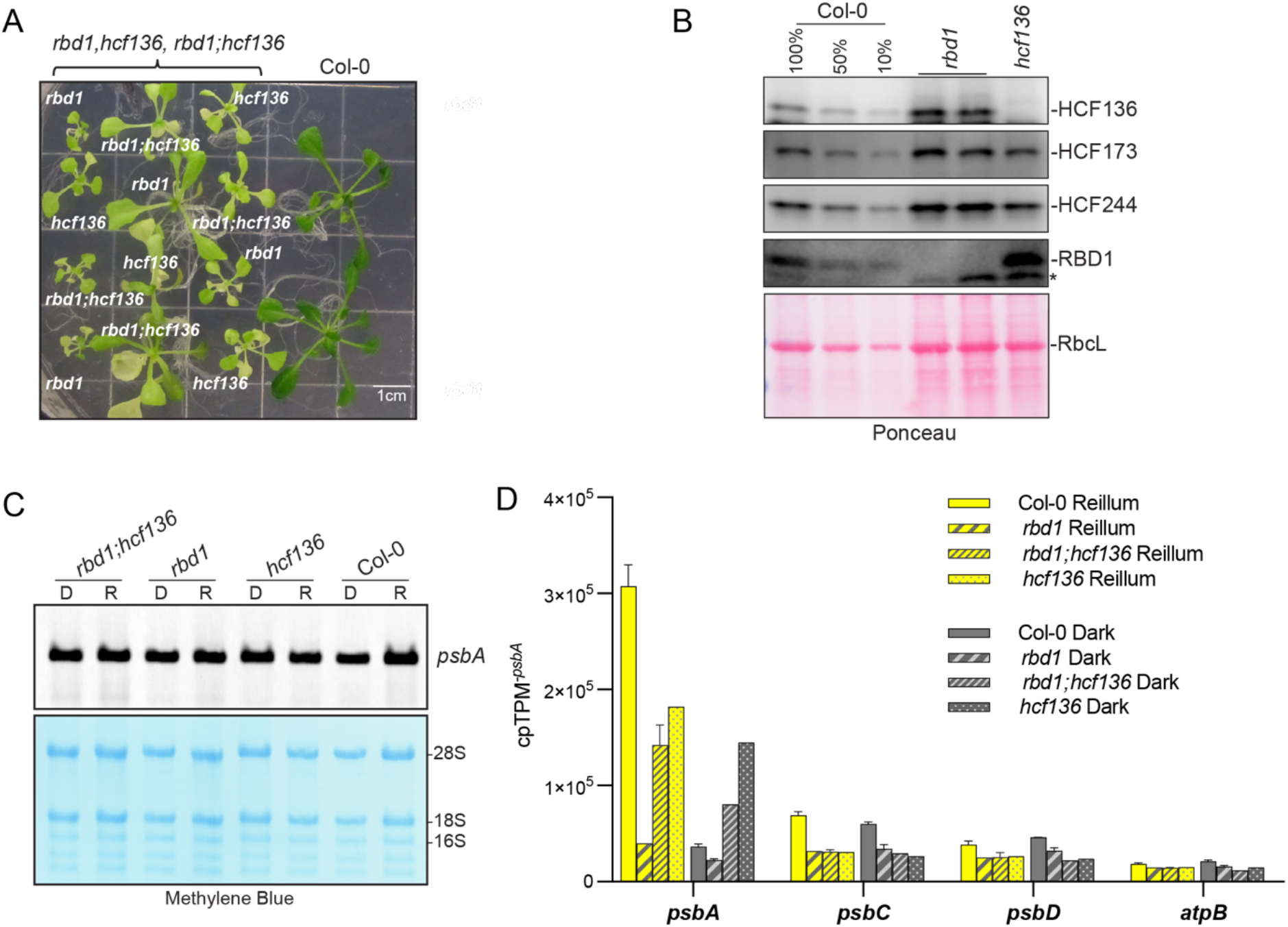
Epistasis analyses using *rbd1;hcf136* double mutants. (A)Phenotypes of single and double mutants. Seeds were germinated and grown on sucrose-containing agar medium. There was considerable heterogeneity in the size of the mutant seedlings of each genotype. The Col-0 seedlings were sown in parallel but grew more rapidly than the mutants. (B) Immunoblot showing the abundance of HCF173, HCF244, HCF136, and RBD1 in *rbd1* and *hcf136* mutants. The same blot was probed sequentially with each antibody, and was stained with Ponceau S to indicate relative sample loading (below). RbcL-Large subunit of Rubisco. *Unknown cross-reacting protein. (C) RNA gel blot analysis of *psbA* mRNA in samples used for ribo-seq in Figure 1 and in Figure 2D. D - 1 h dark adaptation; R - 15 min reillumination. The same blot was stained with methylene blue to visualize rRNAs as a loading control. (D) Ribo-seq analysis of *hcf136;rbd1* double mutants. Plants were harvested after 1-h of dark adaptation or 15-min or reillumination. Graphs show the normalized abundance of ribosome footprints mapping to each indicated chloroplast gene. Data were normalized with the cpTPM^- *psbA*^ method. Values represent the mean +/-SEM of two replicates, with the exception of dark-adapted *rbd1;hcf136* (due to limiting tissue), as well as *hcf136* and reilluminated *rbd1*, for which published datasets corroborate the single replicates presented here (see Fig. 1 and Fig. S1). Data for all chloroplast genes are provided in Dataset S1. The Col-0 and *rbd1* data shown in Figure 1 were extracted from this dataset and are shown again here to facilitate comparisons.

In the ribo-seq analysis, the *hcf136* mutants displayed similarly high *psbA* ribosome occupancy in dark-adapted and reilluminated plants (Fig. 2D). These results were similar to those reported previously (7) (see Fig. S1), except that *psbA* ribosome occupancy in *hcf136* mutants in the current experiment was less than that in illuminated Col-0 but was equivalent to that in illuminated Col-0 in the previous experiments. We suspect that the more advanced developmental stage of Col-0 in comparison to *hcf136* in the current experiment (Fig. 2A) accounts for this difference. Nonetheless, the results from the developmentally-matched mutant samples showed *psbA* ribosome occupancy in illuminated *rbd1;hcf136* double mutants was roughly equivalent to that in *hcf136* single mutants and was much higher than that in *rbd1* single mutants. These results show that RBD1 is not required to stimulate *psbA* translation in the light when HCF136 is missing, indicating that RBD1 activates *psbA* translation in the light by opposing the repressive effect of HCF136.

We were able to obtain only a single replicate of the dark-adapted *rbd1;hcf136* double mutant due to limiting material. Nonetheless, the results suggest that *psbA* ribosome occupancy was considerably higher in dark-adapted *rbd1;hcf136* plants than in dark-adapted *rbd1* or Col-0 plants (Fig. 2D), providing further support for the conclusion that RBD1 activates *psbA* translation by opposing the inhibitory effect of HCF136. Taken together, results above provide strong evidence that RBD1 activates *psbA* translation in the light via light-dependent disruption of a repressive D1 assembly intermediate that requires HCF136.

### HCF244 is required for *psbA* translation in the dark and in the absence of HCF136

Like RBD1, HCF244 is required for *psbA* translation in illuminated seedlings (11, 12) and also promotes an early step in D1’s assembly into the PSII reaction center (2). We investigated how HCF244 affects *psbA* translation as we did for RBD1, asking whether HCF244 is required for *psbA* translation in dark-adapted plants and in the absence of HCF136. We used maize for these experiments because Zm-*hcf244* and Zm-*hcf136* mutants affect *psbA* translation similarly to the orthologous mutants in Arabidopsis (7, 11), and the large maize seed facilitates the recovery of mutant leaf tissue by supporting the rapid growth of non-photosynthetic mutants in soil.

To address whether HCF244 stimulates *psbA* translation in the dark, we performed two independent ribo-seq experiments. One experiment compared *psbA* ribosome occupancy in dark-adapted Zm-*hcf244* mutants and wild-type siblings (Fig. 3, left). The second experiment compared Zm-*hcf244* mutants harvested at midday to those harvested after one hour in the dark (Fig. 3 middle). As a point of comparison, our previously reported results for Zm-*hcf244* and wild-type siblings harvested at midday (11) are shown to the right. The abundance of *psbA* mRNA was previously shown to be unaffected in illuminated Arabidopsis and maize *hcf244* mutants (11, 12), and we confirmed this to be the case in dark-adapted maize seedlings (see Fig. 4C).

**Figure 3.**
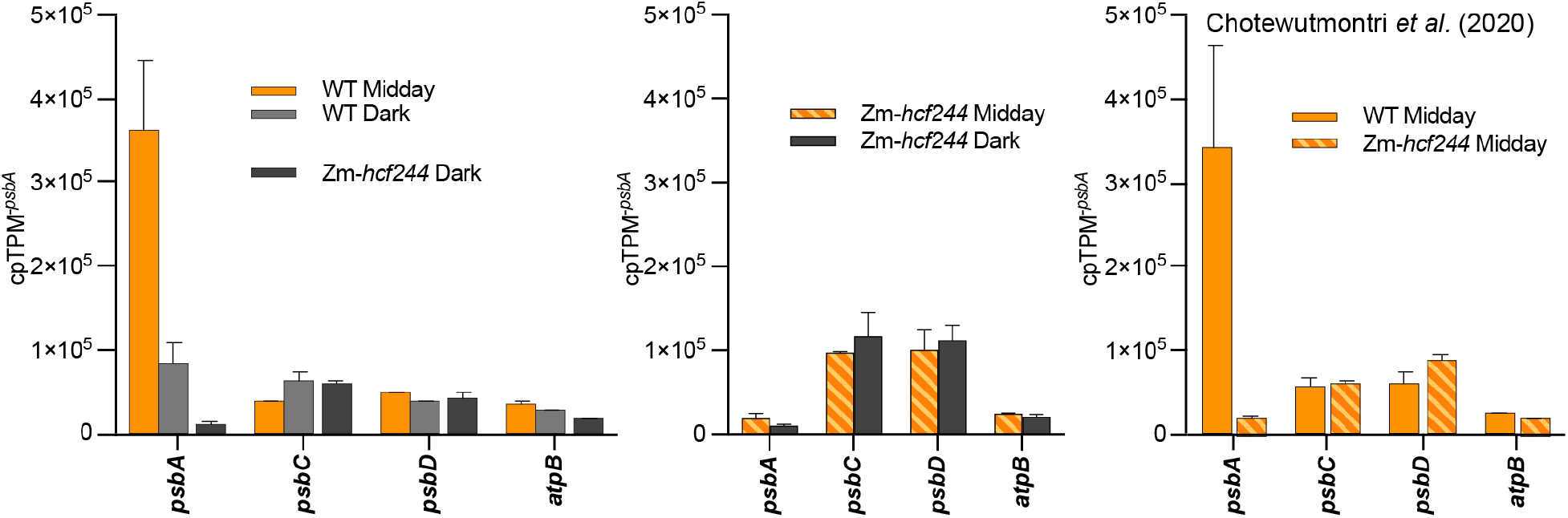
Ribo-seq analysis demonstrating *psbA* translation defect in Zm-*hcf244* mutants in both dark-adapted and illuminated plants. Plants were harvested at midday or after 1-h in the dark. Graphs show the normalized abundance of ribosome footprints mapping to selected chloroplast genes. Data from three independent experiments are shown. Data in the left panel were extracted from the dataset collected in the double mutant analysis presented in Fig. 4. Data in the middle panel come from a comparison of Zm-*hcf244* harvested at midday or after 1-h of dark adaptation. Data in the right panel were reported previously (11). All three datasets were normalized with the cpTPM^-*psbA*^ method. Values are the mean of two replicates +/-SEM. Values for all chloroplast genes are provided in Dataset S1. The abundance of *psbA* mRNA does not change in response to these light shifts or to the presence/absence of Zm-HCF244 (see Fig. 4).

**Figure 4.**
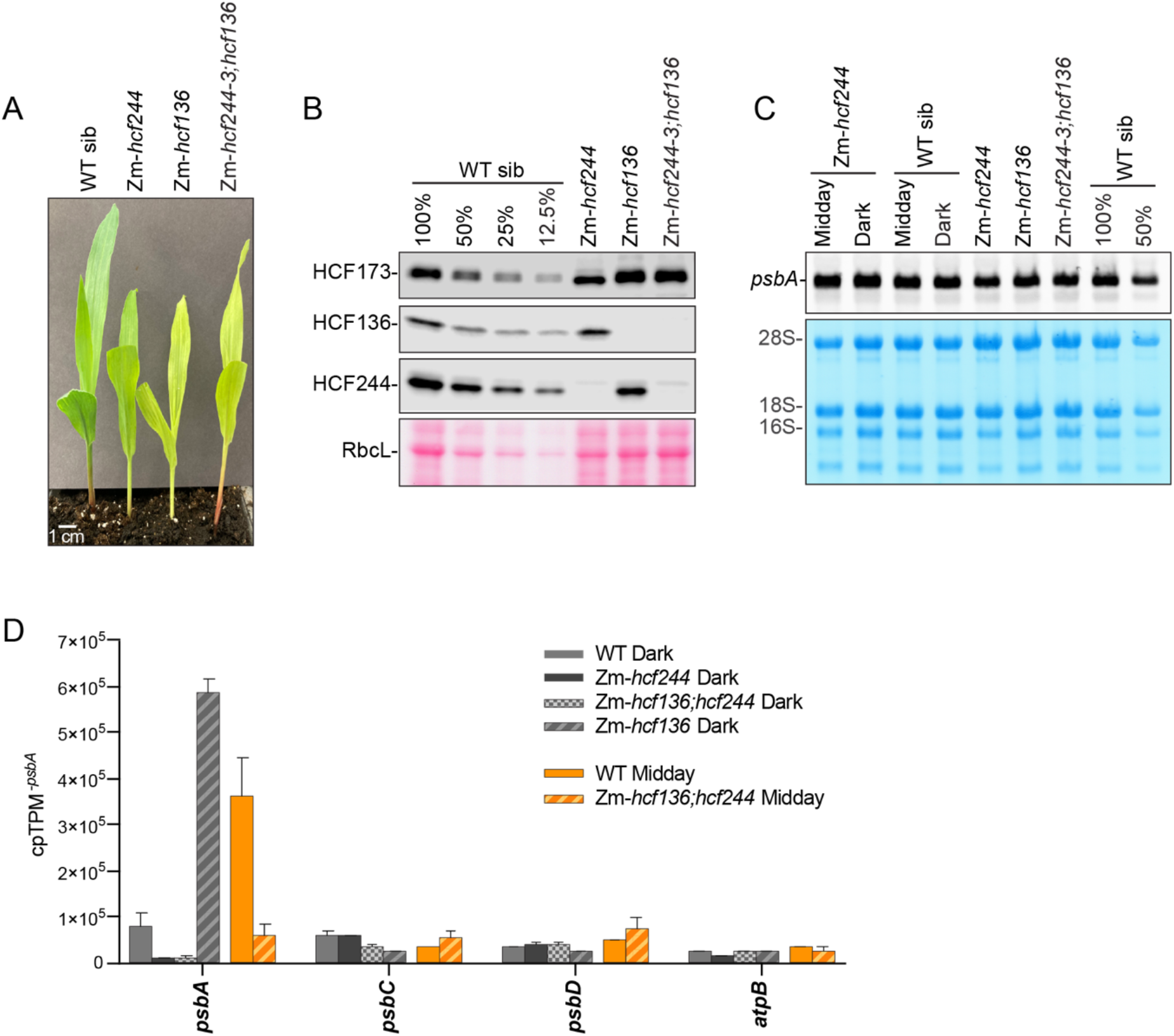
Epistasis analyses using Zm-*hcf244;*Zm-*hcf136* double mutants. (A) Phenotypes of single and double mutants. Seeds were germinated and grown in soil in diurnal cycles for 8 days. (B) Immunoblot showing the abundance of HCF173, HCF244, and HCF136 in plants of the indicated genotypes. The same blot was probed sequentially with antibodies to HCF173 and HCF244, and was stained with Ponceau S to indicate relative sample loading (below). A replicate blot was probed for HCF136. RbcL-Large subunit of Rubisco. (C) RNA gel blot analysis of *psbA* mRNA in samples grown under the same conditions as those used for ribo-seq. RNA was extracted from seedlings harvested at midday or after 1-h of dark adaptation (Dark). The same blot was stained with methylene blue to visualize rRNAs as a loading control. (D) Ribo-seq analysis of Zm-*hcf244;*Zm-*hcf136* double mutants harvested at midday (orange bars) or after 1-h of dark adaptation (black/gray bars). Plants of the indicated genotypes were from the same ear produced by self-pollination of Zm-*hcf244/+*, Zm-*hcf136/+* plants. Data were normalized with the cpTPM^-*psbA*^ method. Values are the mean of two replicates +/-SEM. Data for all chloroplast genes are provided in Dataset S1.The wild-type (dark and midday) and Zm-*hcf244 (*dark) data shown in Figure 3 (left panel) were extracted from this dataset and are shown again here to facilitate comparisons.

The ribo-seq data showed that *psbA* ribosome occupancy in illuminated and dark-adapted Zm-*hcf244* mutants is much lower than that in dark-adapted wild-type plants (Fig. 3). Furthermore, *psbA* ribosome occupancy dropped very little when illuminated Zm-*hcf244* plants were shifted to the dark (Fig. 3 middle). These results show that HCF244 activates *psbA* translation in both the light and the dark, contrasting with RBD1, which activates *psbA* translation by light but has little effect on *psbA* ribosome occupancy in the dark (Fig. 2).

To address whether HCF244 is required for *psbA* translation in the absence of HCF136, we used ribo-seq to analyze *psbA* translation in Zm-*hcf244;Zm-hcf136* double mutants. Progeny of self-pollinated Zm-*hcf244/+*, Zm-*hcf136/+* plants were sown in soil, and homozygous single and double mutants were identified by PCR analysis of small tissue samples. Phenotypes of the single and double mutants are shown in Figure 4A. Immunoblot analysis showed that Zm-HCF173 accumulated to normal levels in all of the mutants, as did Zm-HCF244 and Zm-HCF136 in mutants lacking the other protein (Fig. 4B). The abundance of *psbA* mRNA was similar in all genotypes (Fig. 4C), consistent with prior data (11).

The ribo-seq data showed that *psbA* ribosome occupancy in dark-adapted Zm-*hcf136* mutants was higher than that in illuminated wild-type siblings, consistent with prior results (7) (see Fig. S1) albeit with even more pronounced dysregulation. By contrast, *psbA* ribosome occupancy in dark-adapted Zm-*hcf244*;Zm-*hcf136* double mutants was much lower than that in the wild-type and was similar to that in dark-adapted Zm-*hcf244* mutants (Fig. 4D, black/gray bars). Furthermore, *psbA* ribosome occupancy was much lower in the illuminated double mutant than in illuminated wild-type siblings (orange bars). Together, these results demonstrate that the absence of Zm-HCF136 does not circumvent the need for Zm-HCF244 in either the dark or the light. By contrast, the absence of HCF136 does circumvent the need for RBD1 as a *psbA* translational activator in the light (Fig. 2).

## DISCUSSION

Results reported here elucidate the mechanism underlying the control of *psbA* translation by light and PSII assembly in plant chloroplasts. Prior to this work, it was established that D1 photodamage is the activating signal through which light stimulates *psbA* translation in mature chloroplasts, and that this regulation occurs at the level of translation initiation (6, 7). It was also known that *psbA* translation is disrupted in mutants lacking the PSII assembly factors HCF244 or RBD1, and that it is misregulated in mutants lacking HCF136 (7, 11, 12, 17). Thus, it was clear that there is an intimate connection between D1 assembly, D1 photodamage, and *psbA* translation initiation, but the underlying biochemical connections are not known.

Our findings clarified these connections in several ways. First, we demonstrated that RBD1 and HCF244, which both activate *psbA* translation in illuminated plants, play fundamentally different roles. Whereas HCF244 activates *psbA* translation in both the light and the dark, RBD1 is needed only for the stimulation of *psbA* translation by light. Furthermore, RBD1 and HCF244 exhibit different epistatic relationships with HCF136, which is required to repress *psbA* translation initiation in the dark: whereas HCF244 is required for *psbA* translation even in the absence of HCF136, RBD1 is not. Together these results show that RBD1 functions to counteract HCF136’s repressive effect on *psbA* translation initiation specifically in the light. These findings demonstrate that an antagonistic interplay between HCF136 and RBD1 is at the core of the signal transduction process that senses light-induced D1 damage to modulate *psbA* translation.

The regulatory scheme implied by current data is summarized in Figure 5A. Prior data implicate HCF173 as the direct regulator of *psbA* translation at the end of the signal transduction chain (14, 16, 20, 21). The *psbA* UTRs are sufficient to place a reporter ORF under the control of both HCF173 and HCF244 (20). However, whereas HCF173 binds the *psbA* 5’ UTR (13, 20), there is no reported evidence that HCF244 binds *psbA* mRNA. Results presented here show that HCF244 promotes *psbA* translation via a pathway that is independent of HCF136 and in both the light and the dark. Taken together, these observations suggest that (i) HCF244 acts upstream of HCF173 to facilitate its RNA binding and/or translation activation function, and does so irrespective of D1 photodamage; and (ii) opposing effects of RBD1 and HCF136 act via an intersecting pathway to regulate *psbA* translation in response to D1 photodamage by modulating the activity of HCF173 (Fig. 5A).

**Figure 5.**
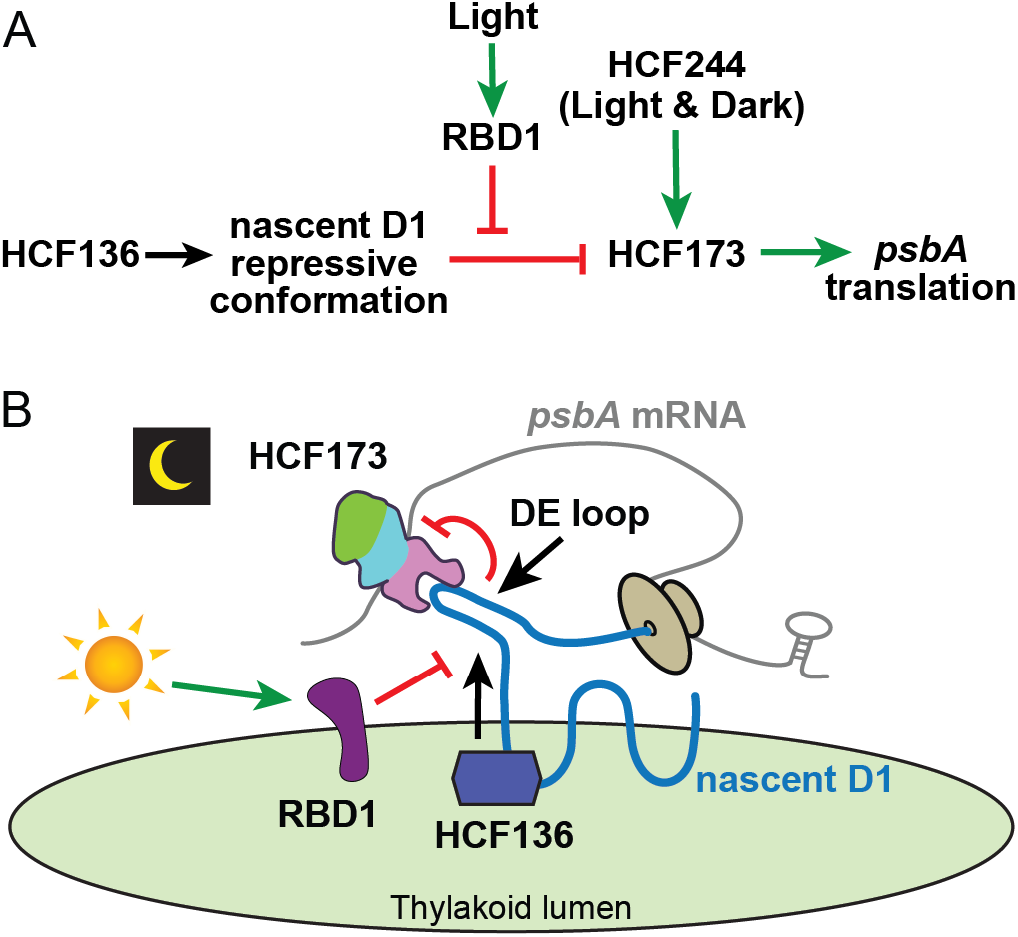
Model for the regulatory circuit connecting HCF136, HCF244, and RBD1 to their effects on *psbA* translation. (A) Regulatory network implied by current data. See Discussion for the rationale for each proposed feature. (B) Working model for the biochemical interactions underlying the effects of RBD1 and HCF136 on light-regulated *psbA* translation. HCF173, which binds the *psbA* 5’ UTR and is required for *psbA* translation (13, 16), harbors a split SDR domain (green and blue) and a CIA30 domain (magenta) (20). HCF136 is proposed to promote a cotranslational interaction between D1’s DE loop and HCF173 that triggers HCF173 to adopt a conformation that does not activate translation; this “OFF” conformation is maintained after translation terminates on the repressed mRNA. Light activates translation via RBD1, which prevents the DE loop from adopting the repressive conformation promoted by HCF136. See Discussion and (20) for the rationale for this model.

How might HCF136 and RBD1 exert their effects on *psbA* translation? HCF136 resides in the thylakoid lumen, so it must affect translation indirectly. The cyanobacterial HCF136 ortholog Ycf48 binds the two lumenal D1 segments that flank its fourth and fifth transmembrane helices (9). There is evidence that this interaction initiates cotranslationally (22), a possibility that is relevant to the regulatory mechanism proposed below. RBD1 is an integral thylakoid membrane protein with a stromal rubredoxin domain (18, 19, 23). Current data suggest that RBD1 acts on D1’s stromal DE loop, possibly in the delivery of a non-heme iron (18). It is intriguing that RBD1 and HCF136 act on the same region of nascent D1 but from opposite sides of the membrane. Thus, an attractive possibility is that they exert their opposing effects on *psbA* translation via their effects on the conformation of D1’s stromal DE loop (Fig. 5B).

The notion that a stromal segment of D1 reports the presence/absence of HCF136 and RBD1 to modulate *psbA* translation is supported by our recent finding that the *psbA* ORF is required in *cis* to repress *psbA* translation in conditions of low D1 photodamage (20). In the same study, we found that a “constitutive” *psbA* translation phenotype is conditioned by a truncated HCF173 mutant, demonstrating that HCF173 is required not only for the activation of *psbA* translation initiation, but also for its repression in the dark. We proposed a mechanism that can account for these newly recognized features of the phenomenology, involving a repressive interaction between nascent D1 emerging from the ribosome and HCF173 bound to the 5’-UTR of the same mRNA molecule (see Fig. 5B) (20). These and other observations culminated in a working model positing that *psbA* translation initiation is repressed in conditions of low D1 damage via a cotranslational interaction between HCF173 and D1’s stromal DE loop, which switches HCF173 to a conformation that does not activate translation but that does remain bound to the *psbA* 5’ UTR. HCF136 represses *psbA* translation in the dark by promoting the repressive conformation of D1’s DE loop (Fig. 5B). The fact that RBD1 influences the conformation of D1’s DE loop (18) and opposes the repressive effect of HCF136 (data shown here) lead us to propose that RBD1 affects the DE loop in a manner that inhibits the DE loop’s interaction with HCF173 (Fig. 5B).

Previously, we proposed an alternative scheme that accounted for the information available at that time (7): we posited (i) that the presence of nascent D1 in the HCF244 assembly complex prevents HCF244 from signaling to HCF173 to activate translation, and (ii) that D1 photodamage relieves that repression by providing a D1-less PSII assembly intermediate into which the HCF244 complex can deposit nascent D1. That model provided an explanation for the constitutively high *psbA* ribosome occupancy in *hcf136* mutants, because HCF136 acts upstream of the HCF244 complex during PSII assembly. However, we no longer favor that model because it does not account for the new observations that the *psbA* ORF represses translation in *cis* and that HCF173 is required for this repression (20).

In summary, results presented here provide evidence that an antagonistic interplay between the PSII assembly factors HCF136 and RBD1 is at the core of the mechanism that modulates *psbA* translation in response to light-induced D1 photodamage. Our data show that light triggers RBD1 to oppose HCF136’s repressive effect on *psbA* translation, and suggest that these effects are mediated by the stromal DE loop of nascent D1 (Fig. 5B). Major outstanding questions include how D1 photodamage changes RBD1’s activity such that it opposes HCF136’s repressive effect, and how HCF244 stimulates *psbA* translation. Although many issues remain to be resolved, it is clear that the mechanism underlying light-regulated *psbA* translation is considerably more intricate than the redox-regulated binding of translational activators to the *psbA* 5’ UTR that has dominated thinking on this topic (24-27). There are, however, intriguing parallels with the “control by epistasy of synthesis” (CES) phenomenon that couples *psbA* translation with D1 assembly in Chlamydomonas (28), whose biochemical basis is unknown. In light of our findings, it may be informative to examine how the HCF136 and RBD1 orthologs influence CES regulation of *psbA* translation in Chlamydomonas.

## MATERIALS AND METHODS

### Plant Material

RBD1 is encoded by Arabidopsis gene AT1G54500. The *rbd1* mutant is T-DNA insertion line SALK_075647. HCF136 is encoded by Arabidopsis gene AT5G23120. The *hcf136* mutant was a generous gift of Joerg Meurer and Peter Westhoff and is described in (10). To generate *rbd1;hcf136* double mutants, plants heterozygous for each mutation were crossed, and double heterozygous progeny were identified by PCR and allowed to self-pollinate. Plants were genotyped by PCR using the primers described in Table S1.

Arabidopsis seeds were sterilized by incubation for 10 min in a solution containing 15% (v/v) bleach and 0.2% (w/v) SDS, followed by a 70% (v/v) ethanol wash. The seeds were then washed three times with sterile water. Seeds were plated and grown in tissue culture dishes containing Murashige and Skoog agar medium (Sigma-Aldrich), supplemented with 3% (w/v) Sucrose. Plants were transferred to fresh growth medium every 7 days. Plants were grown in a growth room in diurnal cycles (16 h of light, 8 h dark 22°C) under Philips F25T8/TL841 Fluorescent Bulbs for 21 d with plates covered with a paper towel (light intensity 40 µmol-m^-2^-s^-1^ under paper towel). One day before harvest, plants were transferred to a different grow room, and illuminated at 40 µE without paper towels (Full light Optoelectronic Materials, LLC. LED Tube light 4500K).

Zm-HCF244 is encoded by maize gene Zm00001eb039850 and Zm-HCF136 is encoded by maize gene Zm00001eb271820. The mutant alleles used here (Zm-*hcf244*-3 and Zm-*hcf136*-1) harbor *Mu-*transposon insertions and were described previously (11). Maize was grown in soil in a growth chamber in diurnal cycles of 16 h light (∼200 µmol-m^-2^-s^-1^ at 28 °C)/8 h dark (26 °C) (Fluorescent bulbs: F72T12/CW/HO and 100 watt incandescent bulbs). To generate Zm-*hcf244*;Zm-*hcf136* double mutants, plants that were heterozygous for each mutation were crossed. Double heterozygous progeny were identified by PCR using the primers listed in Table S1, and were self-pollinated. Progeny that were homozygous mutant for each gene were identified by immunoblotting of small tissue samples of 6-day old seedlings, using antibodies to HCF244 and HCF136.

### Ribosome Profiling

Ribosome footprints were prepared from leaf tissue as described previously (29). For the Arabidopsis experiments, several seedlings were pooled for each replicate and the RNA used for RNA gel blot hybridization was purified from the same lysates. For the maize experiments, ribosome footprints were prepared from the second leaf of 8-day-old seedlings (1 leaf per replicate), at which point the third leaf was beginning to emerge. Experiments were performed with two biological replicates, with the following exceptions: dark-adapted *rbd1;hcf136* double mutant, for which we recovered only enough tissue for one replicate; illuminated *rbd1*, for which the result of our one replicate is corroborated by the published data of Che et al (2022) (see Fig. 1); and Arabidopsis *hcf136* (dark-adapted and reilluminated), for which the results of our one replicate are corroborated by our previous data (7) (see Fig. S1).

Ribo-seq libraries were prepared using the NEXTFLEX Small RNA-Seq Kit v4 (Perkin-Elmer) for all Arabidopsis samples and for the Zm-*hcf244* midday versus 1-h dark experiment shown in Figure 3. The NEXTFLEX Small RNA-Seq Kit v3 was used for the other maize samples. rRNA depletion was performed with the set of custom antisense oligonucleotides described previously (30). The ribo-seq libraries were sequenced by the Genomics and Cell Characterization Core Facility at University of Oregon using Illumina NovaSeq 6000 and NextSeq 2000 in single read 118 bp mode.

Read trimming, alignment and counting were performed as described previously (31). Maize APGv4 assembly and B73 RefGen v4.43 annotation were used for the nuclear and mitochondria genomes, and the chloroplast genome was derived from GenBank accession X86563. Read counts mapping to each chloroplast ORF were normalized for ORF length and sequencing depth with the cpTPM^*-psbA*^ method as follows. Read counts for each chloroplast gene were divided by ORF length (in kilobases) to give reads/kilobase (RPK) for each gene. RPK values were then normalized for sequencing depth by summing RPKs for all chloroplast genes except *psbA* (cpRPK^*-psbA*^) and dividing cpRPK^*-psbA*^ by 10^6^. The RPK value for each gene was divided by cpRPK^-psbA^ /10^6^ to calculate the final normalized value (cpTPM^*-psbA*^). Reads mapping to *psbA* were excluded from the read counts used for normalization because they are so abundant that their inclusion results in a “self-normalization” artifact that reduces the magnitude of genotype- and condition-specific changes in *psbA* translation. Read count and cpTPM^*-psbA*^ values for chloroplast genes are reported in Dataset S1.

### RNA and Protein Blot Analyses

RNA gel blot hybridizations were performed as described previously (32) except that blots were hybridized with a synthetic oligonucleotide probe with an IR800 fluorescent tag (IDT) and were imaged with a Typhoon Imager (Amersham). The *psbA* probe sequence is provided in Table S1. Blots were hybridized overnight in Church Hybridization buffer (7% SDS, 0.5 M NaPhosphate pH 7) at 50°C and washed four times for 5 min each in 1XSSC (0.15 M NaCl, 0.015M NaCitrate) and 0.2% SDS at 50°C.

Proteins were extracted from leaf tissue, fractionated by SDS-PAGE and analyzed by immunoblotting as described previously (32). Antibodies to HCF173 and HCF244 were described in (13) and (11), respectively. The HCF136 and RBD1 antibodies were generous gifts of Peter Westhoff and Joerg Meurer, and Krishna Niyogi, respectively.

## Data Availability Statement

Ribosome profiling data were deposited at NCBI SRA with BioProject number PRJNA1058628.

## Acknowledgements

We are grateful to Kris Niyogi (University of California) for providing antibody to RBD1, and to Peter Westhoff (University of Duesseldorf) and Joerg Meurer (University of Munich) for providing antibody to HCF136. This work was supported by grant MCB-2034758 to A.B. from the National Science Foundation.

## Supporting Information

**Figure S1**. Published ribo-seq data for maize and Arabidopsis *hcf136* mutants (7) harvested at midday, after 1-h of dark-adaptation and 15-min of reillumination.

**Data S1. Summary of ribo-seq data for chloroplast genes**.

## Author Contributions

M.R., P.C., R.W.-C., S.B., and A.B. designed the research, M.R., P.C., R.W.-C., S.B. and E.B. performed the research, M.R., P.C, R.W.-C. and A.B. analyzed the data, A.B. wrote the paper and other authors edited the paper.

---

**Figure S1.**
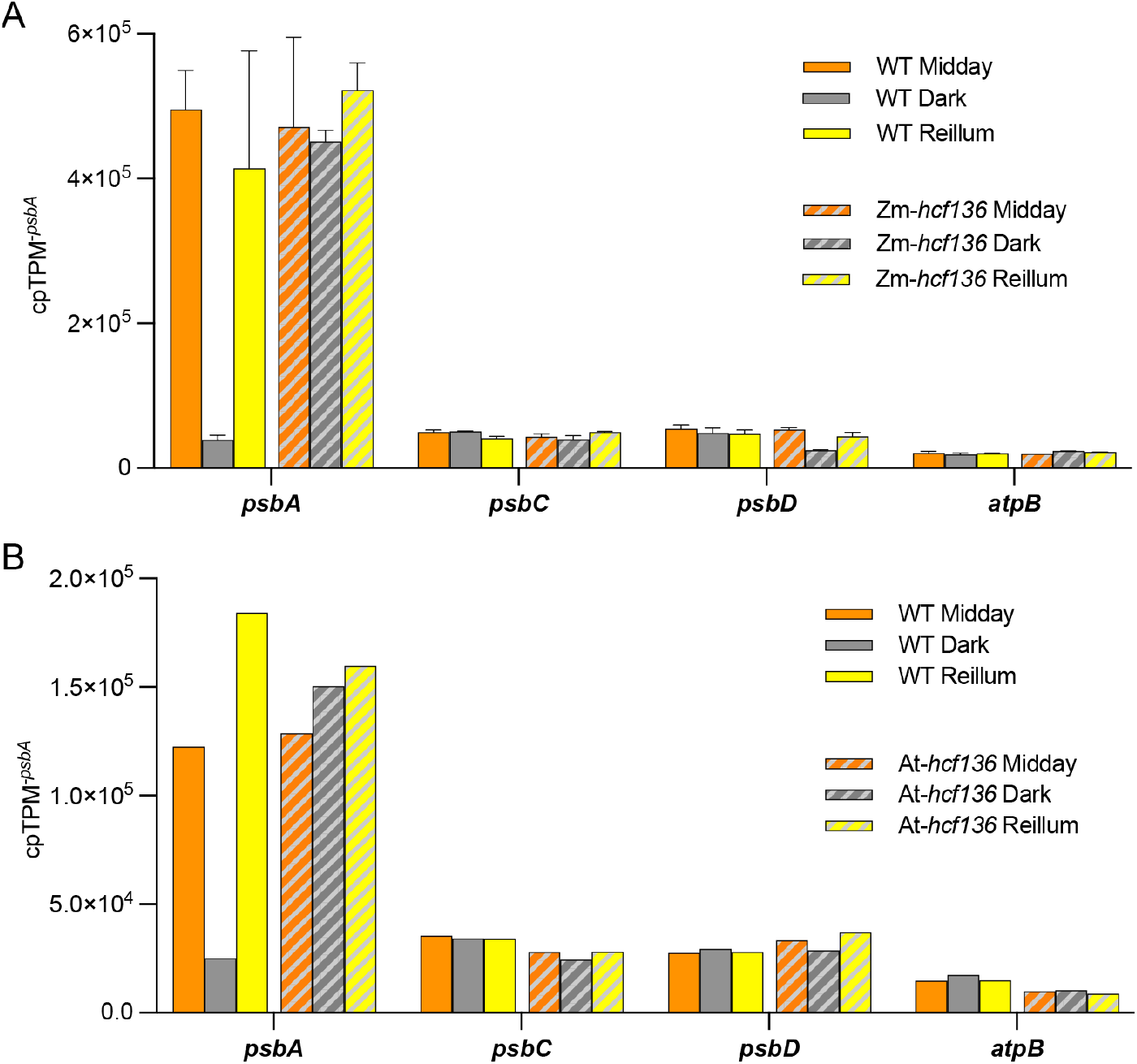
Previously reported ribo-seq data for maize and Arabidopsis *hcf136* mutants (7) harvested at midday, after 1-h of dark-adaptation (Dark) and 15-min of reillumination (Reillum). The data were normalized using the TPM-*psbA* method. Values for the maize experiment (Zm-*hcf136)* are the mean of two biological replicates +/-SEM. The Arabidopsis experiment (At-*hcf136)* was performed with one replicate.

**Table S1.**
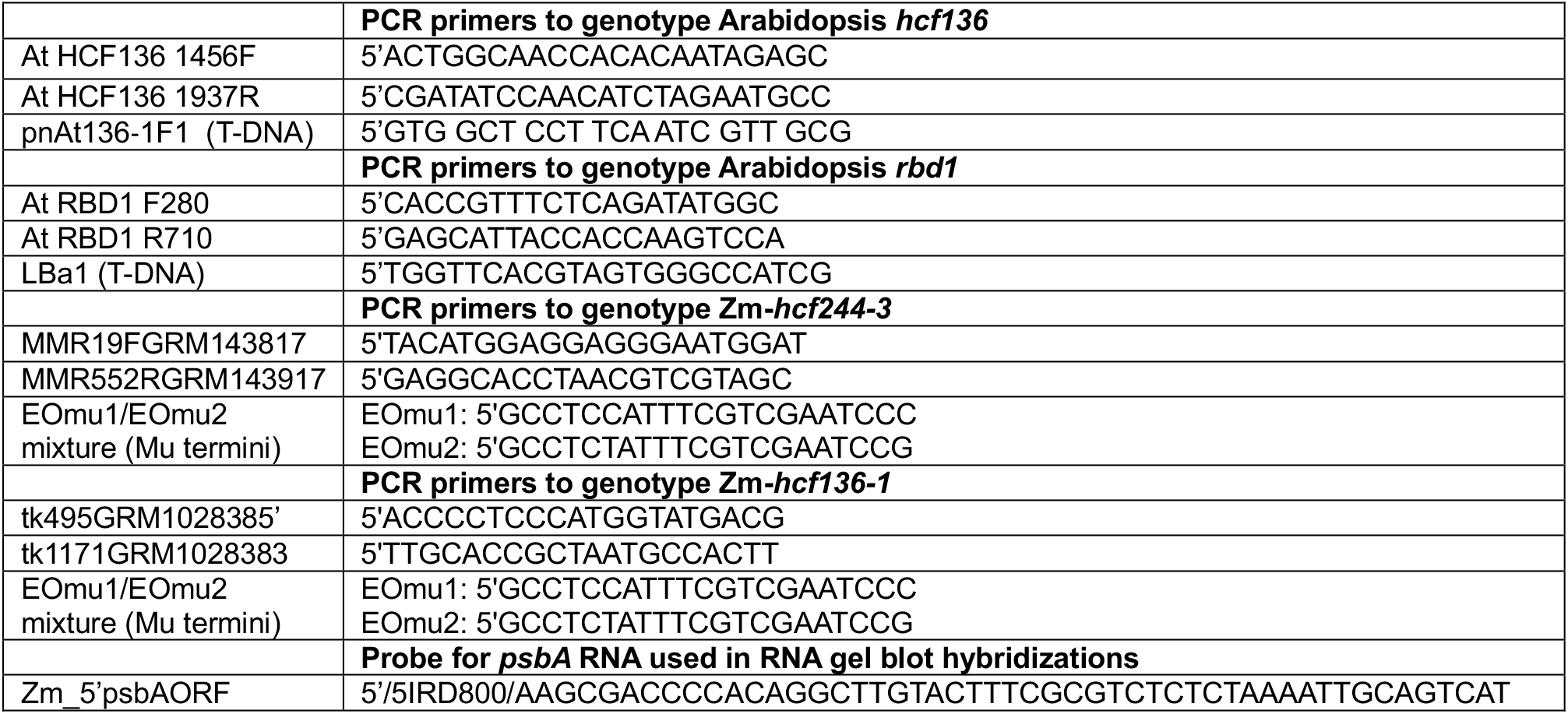
Oligonucleotides used in this study.

